# The effect of local inter-inhibitory connectivity on the dynamics of an activity-dependent neuronal network growth model

**DOI:** 10.1101/052589

**Authors:** Rosanna C. Barnard, Istvan Z. Kiss, Luc Berthouze

## Abstract

The balance between excitation and inhibition in a neuronal network is considered to be an important predictor of neural excitability. Various processes are thought to maintain this balance across a range of stimuli/conditions. However, the developmental formation of this balance remains an open question, especially regarding the interplay between network blue-print (the spatial arrangement of excitatory and inhibitory nodes) and homeostatic processes. In this paper, we use a published model of activity-dependent growth to show that the E/I ratio alone cannot accurately predict system behaviour but rather it is the combination of this ratio and the underlying spatial arrangement of neurones that predict both activity in, and structure of, the resulting network. In particular, we highlight the particular role of clustered inter-inhibitory connectivity. We develop a measure that allows us to determine the relationship between inter-inhibitory connectivity clustering and system behaviour in an exhaustive list of spatial arrangements with a given fixed number of excitatory and inhibitory neurones. Our results reveal that, for a given E/I ratio, networks with high levels of inter-inhibitory clustering are more likely to experience oscillatory behaviour than networks with low levels, and we investigate the network attributes which characterise each global behaviour type produced by the model. We identify possible approaches for extensions of the current work, and discuss the implications these results may have on future modelling studies in this field.

## 1 Introduction

In computational neuroscience, mathematical models of activity-dependent structural plasticity have been used to further our understanding of the development, function, and underlying structure of neuronal networks (Ganguly and Poo, 2013). These models operate on the assumption that network structure results from a number of activity-dependent processes such as neurite outgrowth and growth cone behaviour (Cohan and Kater, 1986; Fields et al., 1990; Schilling et al., 1991), and naturally occurring cell death (Oppenheim, 1991; Ferrer et al., 1992). It has been shown that combining the Shunting model (Grossberg, 1988), describing the neuronal activity of a group of neurons, with terms describing activity-dependent outgrowth, where initially disconnected neurons organise themselves into a network under the influence of endogenous activity, results in a model which can display complex periodic behaviour of electrical activity and network structure, including small-amplitude stable oscillations and intermittent unstable burst-like oscillations (Van Ooyen and Van Pelt, 1996; van Oss and Van Ooyen, 1997). This so-called Extended Shunting model (ES model hereafter) is therefore a useful tool for modelling and analysing the growth and development of neuronal networks, as the qualitative global behaviours reported correspond to brain activity patterns observed in vivo (Buzsaki, 2006).

The balance between excitation and inhibition in a neuronal network is considered to play an important role in determining the cortical computation of the network (Haider et al., 2006), and studies have shown that many human genetic disorders are associated with excessive inhibition in the brain (Ganguly and Poo, 2013). Previous work involving the ES model emphasised the importance of the balance between excitation and inhibition, suggesting that a network developing under conditions of relatively high inhibition is more likely to end up in a ‘pathological’ state with oscillatory behaviour, whereas a network developing under a low level of inhibition during its development enables the system to move to the ‘normal’ state, where the network structure reaches a stable end state with stable levels of activity (van Oss and Van Ooyen, 1997). The oscillatory behaviour induced by moderate proportions of inhibition was thought to be a result of interactions between excitatory and inhibitory activity, occurring on the timescale of the membrane potential (electrical activity) (Van Ooyen et al., 1995).

Central to this paper is another factor which has been identified by Van Ooyen et al. (1995) as affecting system behaviour beyond the proportion of inhibitory neurons, namely, the spatial arrangement of inhibitory neurons. Whilst inhibitory neurons induce outgrowth and allow disconnected subnetworks to become connected, the distribution of inhibitory neurons plays an important role in the interplay between structure and function. When highly-clustered inhibitory cells (with self-inhibition) become electrically inhibited, their outgrowth is stimulated, inducing long-range inhibition. This in turn means that these networks develop stronger excitatory-excitatory, excitatory-inhibitory (and inhibitory-excitatory) connectivity.

Still, there is a lack of a comprehensive account of the extent to which the spatial organisation of excitatory and inhibitory neurons influences global system behaviour and the structural characteristics of the resulting neuronal networks. Such an account is made necessary by the a growing body of evidence suggesting the presence of non-random network attributes within neuronal network structures, including modular community structure, highly connected and central hub regions, short characteristic path length (associated with high global efficiency of information transfer), high levels of clustering (associated with robustness to random error), and degree distributions compatible with the existence of highly-connected hub neurons (Bullmore and Sporns, 2009; Sporns, 2011; van den Heuvel and Sporns, 2013). In this paper, we investigate how much influence the spatial arrangement of excitatory and inhibitory neurons, and more specifically, inhibitory clustering, has on the global system dynamics of network structure and electrical activity in the ES model.

In what follows, we begin by outlining the differential equations governing the activity-dependent neuronal network growth model, giving a detailed description of each model element, and explaining how these elements combine to create a system in which each and every neuron strives to maintain a desired level of electrical activity. We explain the choice of model parameters, describe the implementation of the growth simulations, and outline the terminologies and notation used throughout this paper. In Section 3, we show that the proportion of inhibition in any system of neurons alone is not a sufficient, but a necessary condition for inducing oscillatory behaviour. We define a method for measuring the extent of inhibitory clustering within our networks, and demonstrate that higher levels of inhibitory clustering induce oscillatory behaviours, whilst lower levels of inhibitory clustering lead the networks to a final, stable end state. Finally, we look at the qualitatively distinct global network behaviours produced by the model, and attempt to understand what characterises each behaviour in terms of network theoretical measures. We conclude with a discussion of our findings and their possible implications with regards to future work in this field.

## 2 Methods

### 2.1 Model

This work is based on the model of neuronal development proposed by Van Ooyen et al. (1995). It extends the Shunting model proposed by Grossberg (1988) by incorporating activity-dependent outgrowth terms. This allows a network containing excitatory and inhibitory neurons to be grown from initial conditions. Each neuron is assigned a unique spatial location (see Section 2.2.2) and modelled with a variable radius, denoting the area of its influence. When the circular fields of two neurons overlap, those neurons become connected with a strength proportional to the area of overlap, multiplied by the appropriate synaptic efficacy term, determined by the type (excitatory or inhibitory) of the two neurons.

Precisely, the system is governed by the following dimensionless coupled differential equations describing the rate of change of electrical activity and neuritic radii of N excitatory and *M* inhibitory neurons denoted X and *Y* respectively:

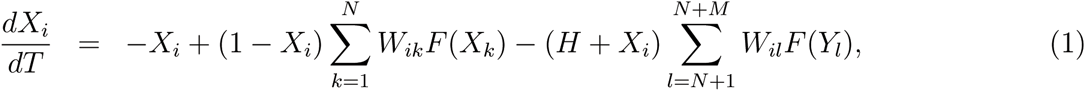

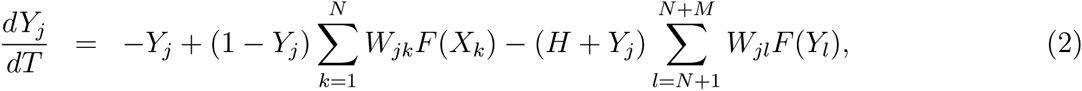

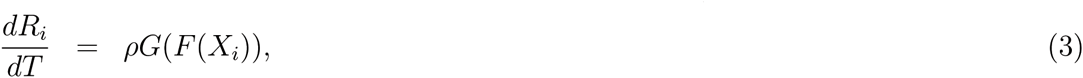

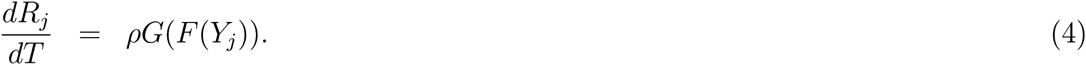

In the first two equations, which correspond to the original Shunting model, *X*_*i*_ and *Y*_*j*_ can be referred to as membrane potentials of excitatory neuron *i* ∈ {1,…, *N*} and inhibitory neuron *j* ∈ {*N* + 1,…, *L* = *N*+ *M*}, respectively, expressed in units of saturation potential (finite maximum). Membrane potentials have a finite minimum that is — *H* times the saturation potential. A change in membrane potential of a neuron is influenced by its own electrical activity and that of any other neuron it shares a connection with. The strength of the connection between neurons *i* and *j* at any point in time is given by element *W*_*ij*_ of the (symmetric) weight matrix ***W***. In this weight matrix, self-overlap terms are set to zero (*W*_*ii*_ = 0 ∀ *i* ∈ {1,…,*N* + *M*}) and non-diagonal elements are calculated as

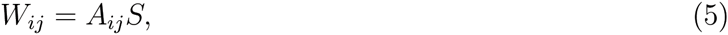
where *A*_*ij*_ describes the area of overlap between the circular neuritic fields of neurons *i* and *j*, and *S* describes a synaptic strength parameter dependent on the type (excitatory or inhibitory) of the two neurons *i* and *j*.

Equations (3) and (4) result from extending the Shunting model to incorporate activity dependency. *R*_*i*_ and *R*_*j*_ denote the radii of the neuritic field of excitatory neuron *i* ∈ {1,…, *N*} and inhibitory neuron *j* ∈ {1,…, *M*}, respectively. Parameter *ρ* describes the rate of outgrowth of the neuritic radii (*ρ* is constant in our analyses) and the functions *F* and *G* are continuous saturating functions:

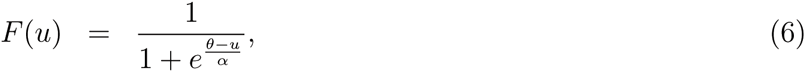

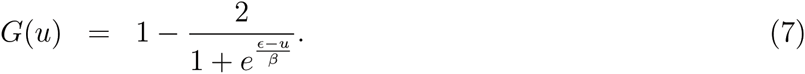

The function *F* can be thought of as governing the mean firing rate of each neuron, with values in the bounded range (0,1). The function *G* describes the growth rate of a neuron as a function of the neuron’s firing rate, and has values in the range (-1,1). *θ*, *α*, *∊* and *β* are parameters whose role and default values are given in Table 1. The value of e has great influence on system behaviour, since each neuron attempts to maintain a membrane potential value equal to ∊. When a neuron’s membrane potential stabilises at ∊, the growth function *G* tends to zero and the neuritic radius will cease to change. If all neurons in the system are able to maintain a membrane potential equal to ∊ simultaneously, a system steady state is reached.

**Table 1.** Default values for all model parameters used in simulations

| Parameter name | Parameter description | Default parameter value |
| --- | --- | --- |
| $\rho$ | rate of growth of neuritic field | 0.0001 |
| $\theta$ | firing threshold for firing rate function $F$ | 0.5 |
| $\alpha$ | gradient of firing rate function $F$ | 0.1 |
| $\beta$ | gradient of growth function $G$ | 0.1 |
| $\epsilon$ | threshold for growth function $G$ | 0.6 |
| $H$ | ratio between minimum and maximum potentials | 0.1 |
| $S_{ee}$ | excitatory to excitatory synaptic efficacy | 1 |
| $S_{ie}$ | excitatory to inhibitory synaptic efficacy | 1 |
| $S_{ei}$ | inhibitory to excitatory synaptic efficacy | 1 |
| $S_{ii}$ | inhibitory to inhibitory synaptic efficacy | 1 |

### 2.2 Model implementation

All simulations were implemented in the Matlab environment, using a variable-order stiff ode solver *(ode15s)*. Initial conditions were specified, consisting of the spatial location of all neurons within the system and the value of each neuron’s radius and membrane potential at time zero. In all our experiments, the neurons were placed on a one-dimensional lattice with periodic boundary conditions. The distance between neurons was nominally set to 1. The spatial arrangement of excitatory and inhibitory neurons is detailed in Section 2.2.2. Initial membrane potential values were set to zero (the resting potential level) for all neurons. Different scenarios of initial radii values were used as specified in each of the relevant sections. These included:

- randomly (uniformly) generated values in the range (0, 0.6) creating an initially largely disconnected network,
- identical values (radii=0.4) creating a totally disconnected network,
- identical values (radii=0.6) for a regular ring network.

We imposed that all neuritic radii be greater or equal than zero at all times. Simulations were run until system behaviour could be clearly identified (typically up to time > 10^5^).

### 2.2.1 Parameter choice

Our analyses used similar parameters to those used by Van Ooyen et al. (1995) and are summarised in Table 1. They differed in two respects. First, whereas Van Ooyen et al. set the proportion of inhibitory neurons in the range of 10-20%, in accordance with in vivo observations in various brain regions (Van Ooyen et al., 1995), we systematically varied this proportion in order to provide a fuller picture of system behaviour in relation to the balance between excitation and inhibition (see also the next Subsection). Second, for simplicity, we kept the synaptic efficacy parameters (*S*_*ee*_, *S*_*ie*_, *S*_*ei*_ and *S*_*ii*_) identical throughout.

### 2.2.2 Spatial arrangement of excitatory and inhibitory neurons

Since connectivity is determined by the amount of overlap between neuritic fields centred over fixed neurons, it follows that the joint degree distributions for X (excitatory) and Y (inhibitory) neurons will strongly depend on the spatial arrangement of these neurons, and particularly, whether they are well mixed or clustered – an operational definition of this clustering will be given in Section 3.2. We used a systematic approach to building arrangements of excitatory and inhibitory neurons assigned to lattice positions. We define a *tile* as a single row of *l* ∈ ℤ^+^ neurons, with nominal distance 1 between consecutive neurons. Then, complex one-dimensional networks with periodic boundary conditions could be formed by duplicating tiles a specified number of times. For example, a network of size *L* = 12 with a 1/3 proportion of inhibitory neurons could consist of a single tile of length *l* = 12 with 4 inhibitory neurons or 4 identical tiles of length *l* = 4 with 1 inhibitory neuron each (see example below). The use of periodic boundary conditions means that any pair of neurons in the network will be at most 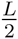 =(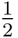Xnumber of neurons in the network) distance away from one another. When representing spatial arrangements throughout this paper, we use filled circles to denote inhibitory neurons, and empty circles to denote excitatory neurons, e.g., •∘∘•∘∘•∘∘•.

## 3 Results

### 3.1 The proportion of inhibitory neurons - a necessary condition for global system behaviour

In our analyses it was found that the proportion of inhibition alone cannot accurately predict the system’s global behaviour. Simulations with tile size 13 were implemented where the proportion of inhibition in the system was set to 9/13 ≈ 30.77% with all other parameters held constant, bar the placement of neurons. For the initial conditions, all membrane potential values were set to zero (the resting potential), and all neuritic radii values were set to 0.4, creating a wholly disconnected network. A number of spatial arrangements were considered, randomly picked from a set of unique tile arrangements generated by a Matlab script that selected at random *M* out of *L* positions for inhibitory neurones, and placed *N* excitatory neurones onto the remaining lattice positions (note that a more systematic approach is used in Section 3.2).

Three qualitatively distinct global behaviours are found (Fig. 1), namely, (a) stable periodic oscillations of small amplitude of both network structure and electrical activity (Figs. 1a and 1b), (b) large amplitude unstable oscillations in the network structure and unstable burst-like oscillations in electrical activity (Figs. 1c and 1d), and (c) complete stabilisation of both network structure and electrical activity (Figs. 1e and 1f), all occurring after an initial period of overshoot and pruning of connections.

**Figure 1:**
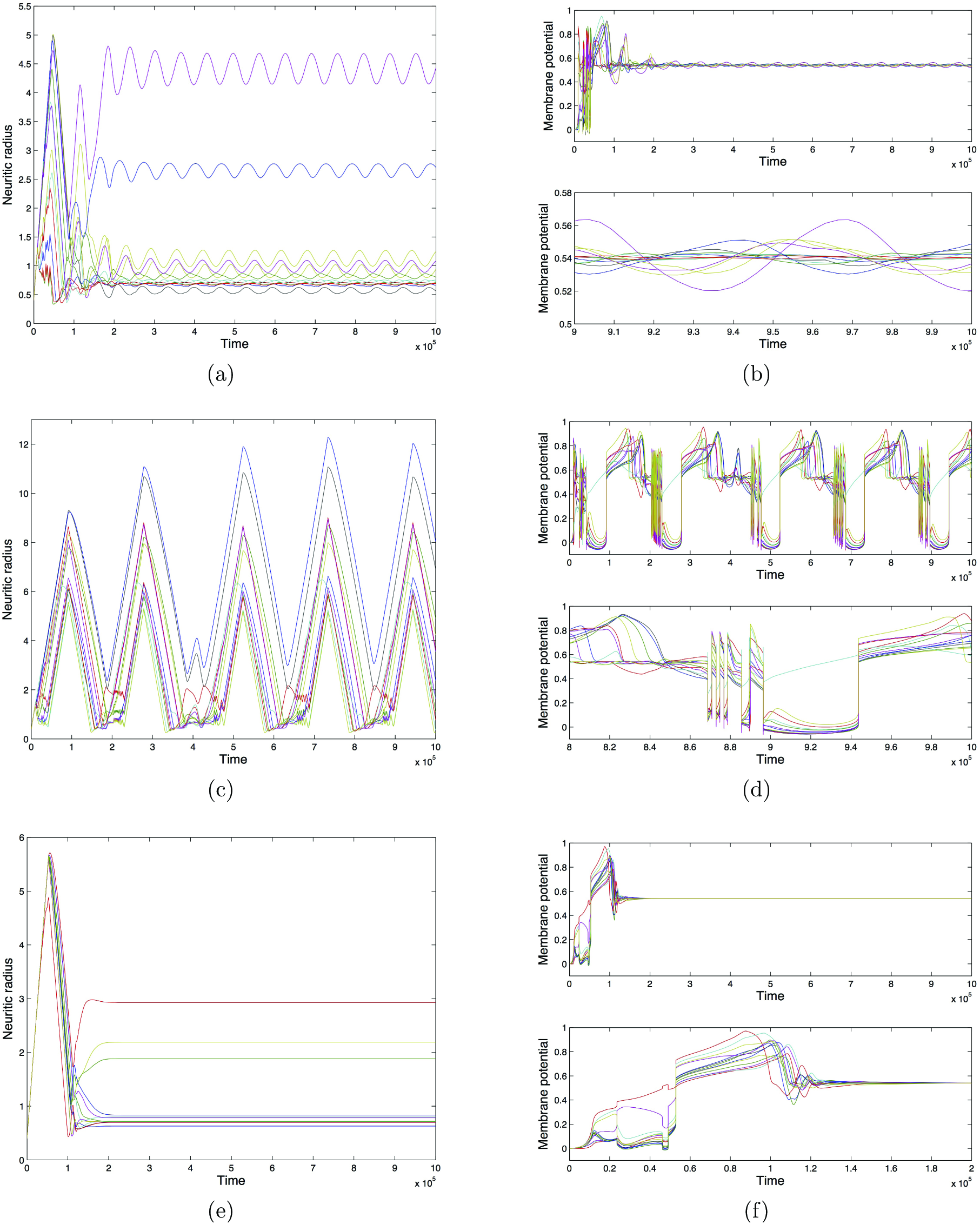
Global system behaviour of mixed 1D lattice networks with various spatial arrangements of neurons. In all cases L = 13, *N* = 9, M = 4 and default parameters listed in Table 1 are used. The networks are initially wholly disconnected and neurons are at rest. Figures on the left-hand side depict the dynamics of the individual neuritic radii values; figures on the right-hand side depict the dynamics of the individual membrane potential values. Neuron arrangements are as follows: ∘ ∘ ∘ ∘ ∘ • • ∘ ∘ • ∘ • ∘ (a,b), ∘ ∘ • ∘ ∘∘ ∘∘ • • • ∘ ∘ (c,d), ∘ ∘ • ∘ • ∘ ∘ • ∘ ∘∘ • ∘ (e,f).

Since all networks were initialised with identical neuritic field sizes and membrane potentials, we conclude that varying the spatial arrangement of neurons within the network can induce distinct global behaviours. Therefore, the proportion of inhibition alone may be a necessary but not *sufficient* condition to induce oscillatory system behaviour.

### 3.2 The effect of inhibitory clustering on global system behaviour

The occurrence of different global behaviours when the proportion of inhibitory neurons in a tile is fixed and simulations of various spatial arrangements are run (Fig. 1) suggests that some feature of spatial organisation of neurons (and specifically inhibitory neurons) could impact the presence of oscillatory behaviour. As most models typically assume a well-mixed population, here, we focus on the possibility that inter-inhibitory connectivity induced by spatial clustering of inhibitory neurons could be an important determinant. Measuring such spatial clustering is not straightforward and we are not aware of any established method for quantifying it. In what follows, we define the measure of inhibitory clustering in a tile network structure with periodic boundary conditions as

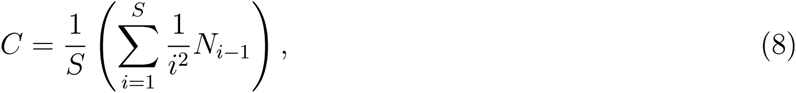
where *S* is the shortest distance between the first and last inhibitory neurons in a tile, taking periodic boundary conditions into account, and *N*_*i*_ denotes the number of pairs of inhibitory neurons with *i* excitatory neurons separating them. *N*_*i*_ is consistently counted without multiplicity, meaning that the total *N*_*i*_ count will always be equal to (*M* — 1) with *M* inhibitory neurons, and any *N*_*i*_ pairs of inhibitory neurons should not have another inhibitory neuron in between them. For example, network ••∘∘•∘∘∘∘∘ has *S* = 4, *N*_*0*_ = 1 (the only adjacent pair of inhibitory neurons), *N*_*2*_ = 1 (the single pair of inhibitory neurons separated by 2 excitatory neurons), and *N*1 = *N*3 = *N*4 = 0 leading to a clustering value 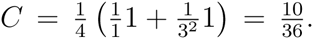 The inhibitory clustering measure takes value 1 when the greatest possible level of inhibitory clustering is achieved in a tile, i.e., when all inhibitory neurons are adjacent to one another and hence *N*_*0*_ = *S*, and its value tends towards zero as the extent of inhibitory clustering decreases. The decay of the measure is controlled by the power of the weighing factor. This power was selected as a trade-off between an insufficient penalty for the spreading out of inhibitory neurons (for example, 1/*i*) and insensitivity to subtle variations in the structure of the tile (for example, 1/*i*^3^).

Simulations with fixed tile size and number of inhibitory neurons, but various levels of inhibitory clustering, revealed distinct global behaviours and a trend suggesting that networks with a higher clustering measure are more likely to experience oscillatory behaviours (Table 2). In networks of size eight containing two inhibitory neurons (Fig. 2), a wider range of final neuritic field sizes were observed as the spatial proximity of the inhibitory neurons was increased, and once the inhibitory neurons became immediate neighbours, periodic oscillatory behaviour occurred in both the network structure (neuritic radii values) and the electrical activity of the network itself (Figs. 2a and 2b). Here, an idiosyncratic behaviour of the model is observed whereby the largest neuritic field in the network continues to grow whilst experiencing stable oscillations (Fig. 2a). This behaviour results from the fact that once the neuron has become completely connected with every other neurons in the network, further growth has no effect on the connection strengths to itself and others and outgrowth continues unbounded.

**Figure 2:**
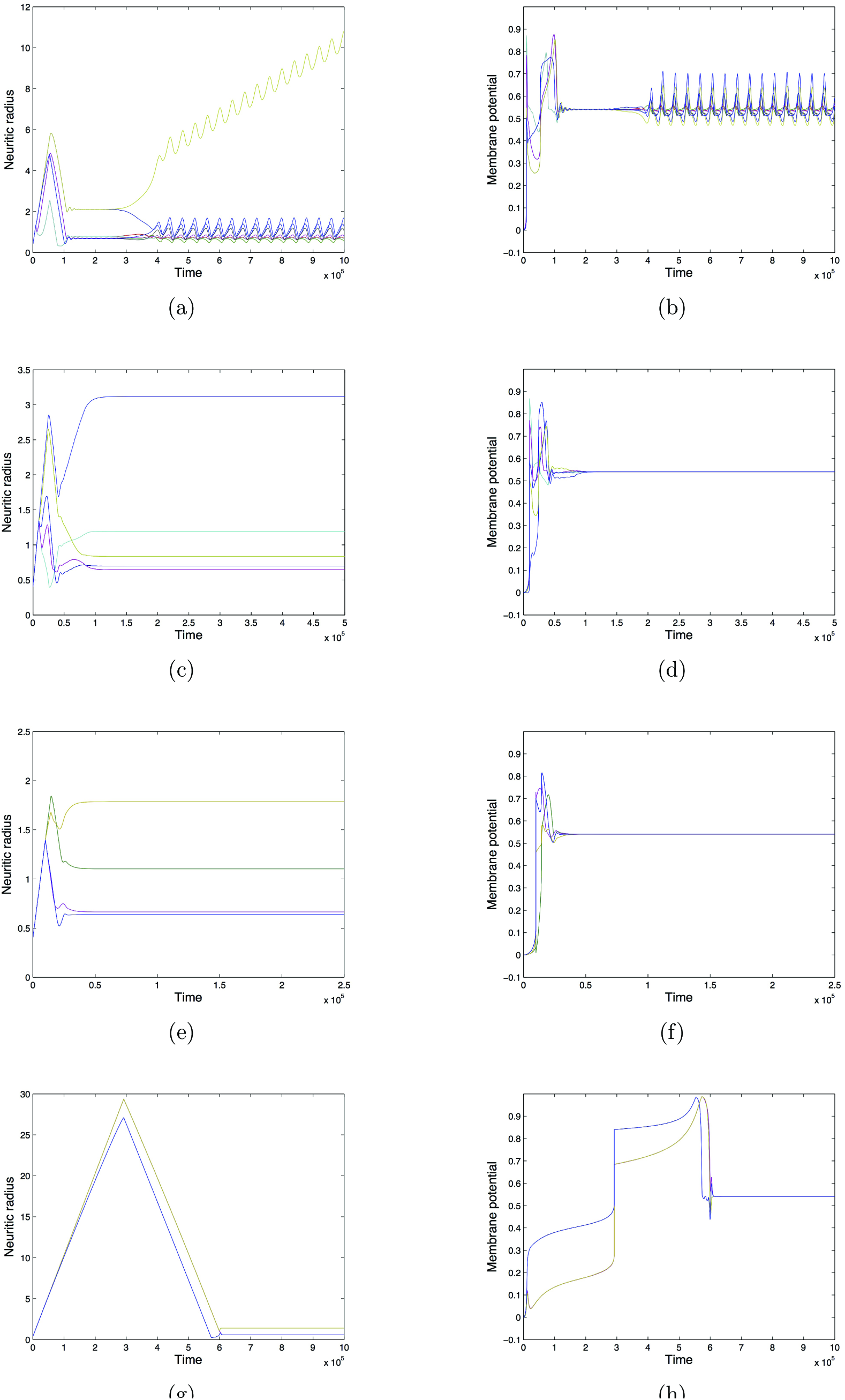
Global system behaviour of mixed 1D lattice networks with various levels of inhibitory clustering. In all cases L = 8, *N* = 6,*M* = 2 and default parameters listed in Table 1 are used. The networks are initially wholly disconnected and neurons are at rest. Figures on the left-hand side depict the dynamics of the individual neuritic radii values; figures on the right-hand side depict the dynamics of the individual membrane potential values. Neuron arrangements are as follows: • • ∘ ∘ ∘ ∘ ∘∘ (a,b), • ∘ • ∘ ∘ ∘ ∘∘ (c,d), • ∘ ∘ • ∘ ∘ ∘∘ (e,f), • ∘ ∘ ∘ • ∘ ∘∘ (g,h).

**Table 2.**
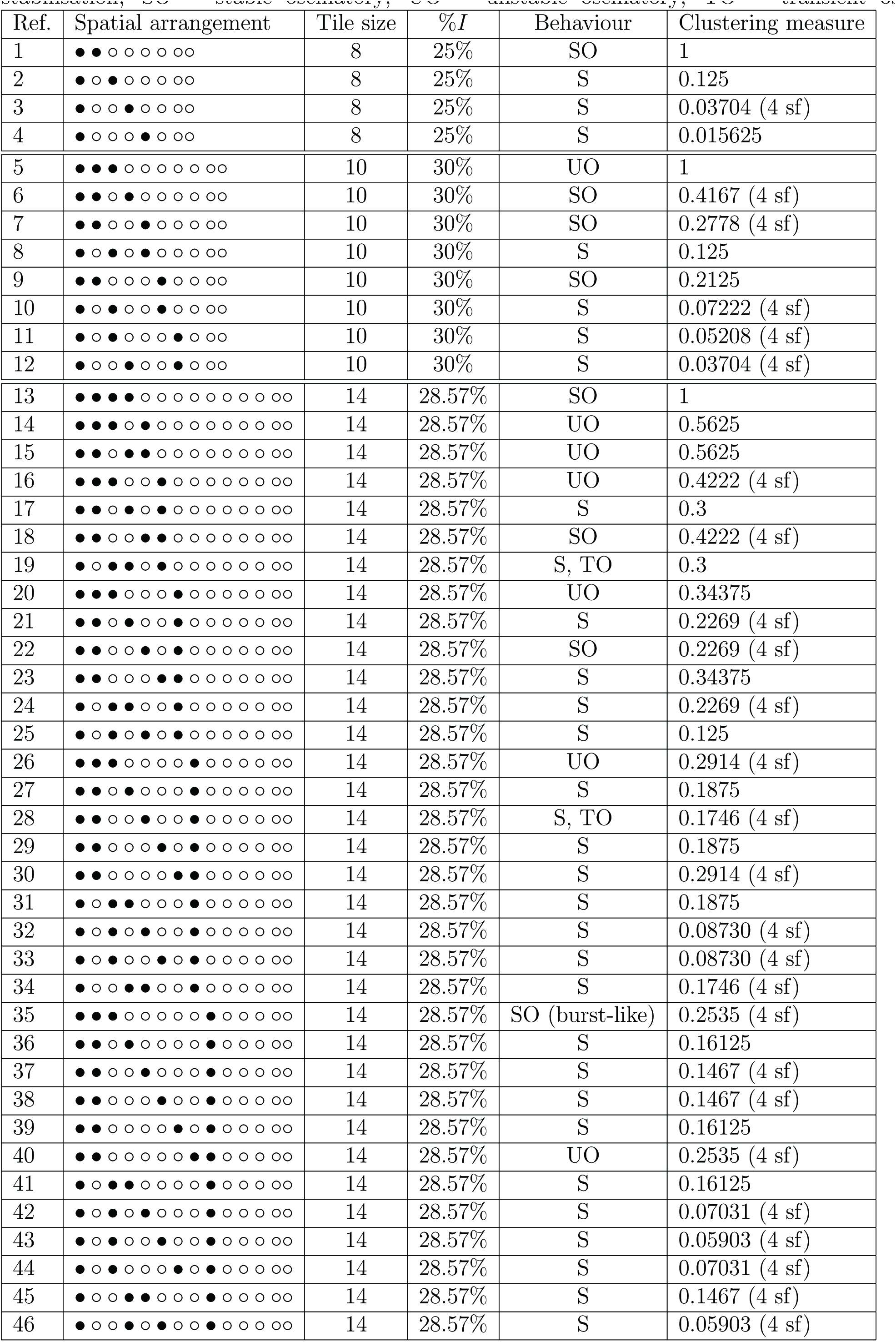
Global system behaviour and clustering measure for mixed 1*D* simulations with various spatial arrangements. Default values were used on initially disconnected networks. S = stabilisation; SO = stable oscillatory; UO = unstable oscillatory; TO = transient oscillations.

In networks containing three inhibitory neurons and seven excitatory neurons (Fig. 3), the most extreme oscillatory behaviour was again seen when the inhibitory clustering is at its greatest (Figs. 3a and 3b). In this case, the network structure and membrane potentials experienced a bout of periodic oscillations, before developing unstable burst-like oscillations of much greater amplitude. As the extent of inhibitory clustering decreases slightly, the system experiences stable oscillations with a smaller amplitude, in both network structure and electrical activity (Figs. 3c and 3d). Eventually, the inhibitory neurons become more evenly distributed across the lattice positions and the neuritic radii and their associated electrical activity experience complete stabilisation (Figs. 3e - 3h). An exhaustive list (taking symmetries and reflections into account) of all possible spatial configurations for tile sizes eight, ten and fourteen, containing two, three and four inhibitory neurons respectively, also suggests that higher inhibitory clustering increases the likelihood of oscillatory behaviour being observed (Table 2).

**Figure 3:**
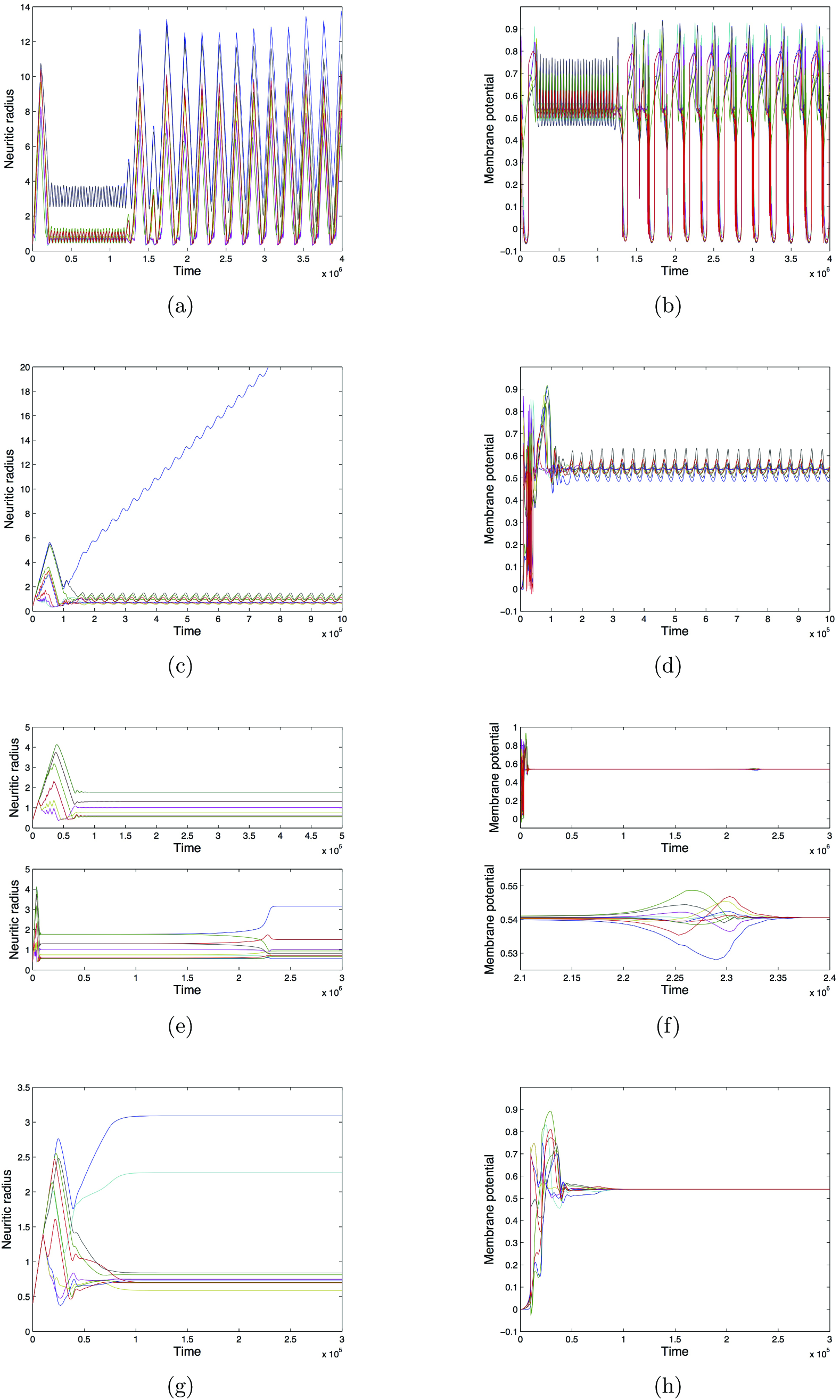
Global system behaviour of mixed 1D lattice networks with various levels of inhibitory clustering. In all cases*L* = 10,*N* = 7, *M* = 3 and default parameters listed in Table 1 are used. The networks are initially wholly disconnected and neurons are at rest. Figures on the left-hand side depict the dynamics of the individual neuritic radii values; figures on the right-hand side depict the dynamics of the individual membrane potential values. Neuron arrangements are as follows: •••∘∘∘∘∘∘∘ (a,b), • • ∘ • ∘ ∘ ∘ ∘ ∘∘ (c,d), • ∘ • ∘ • ∘ ∘ ∘ ∘∘ (e,f), • ∘ • ∘ ∘ • ∘ ∘ ∘∘ (g,h).

A systematic mapping of the relationship between inhibitory clustering and the proportion of inhibitory neurons reveal the emergence of distinct regimes (Fig. 4). There are no significant differences for the regimes observed between simulations with initially disconnected networks (radii=0.4, Fig. 4a) and those starting with a regular ring network (radii=0.6, Fig. 4b). Networks with lower proportions of inhibitory neurons, and lower clustering values are more likely to experience complete stabilisation of electrical activity and network structure. However, a handful of configurations with *high* inhibitory clustering and low proportions of inhibitory neurons also experienced complete stabilisation. In general, as inhibitory clustering increases, systems developed oscillatory behaviours. We observe that the oscillatory regime also extends into slightly higher proportions of inhibitory neurons than the stabilisation regime. Past the oscillatory regime, with the highest proportions of inhibitory neurons, and a range of inhibitory clustering values, we see systems experience unbounded growth.

**Figure 4:**
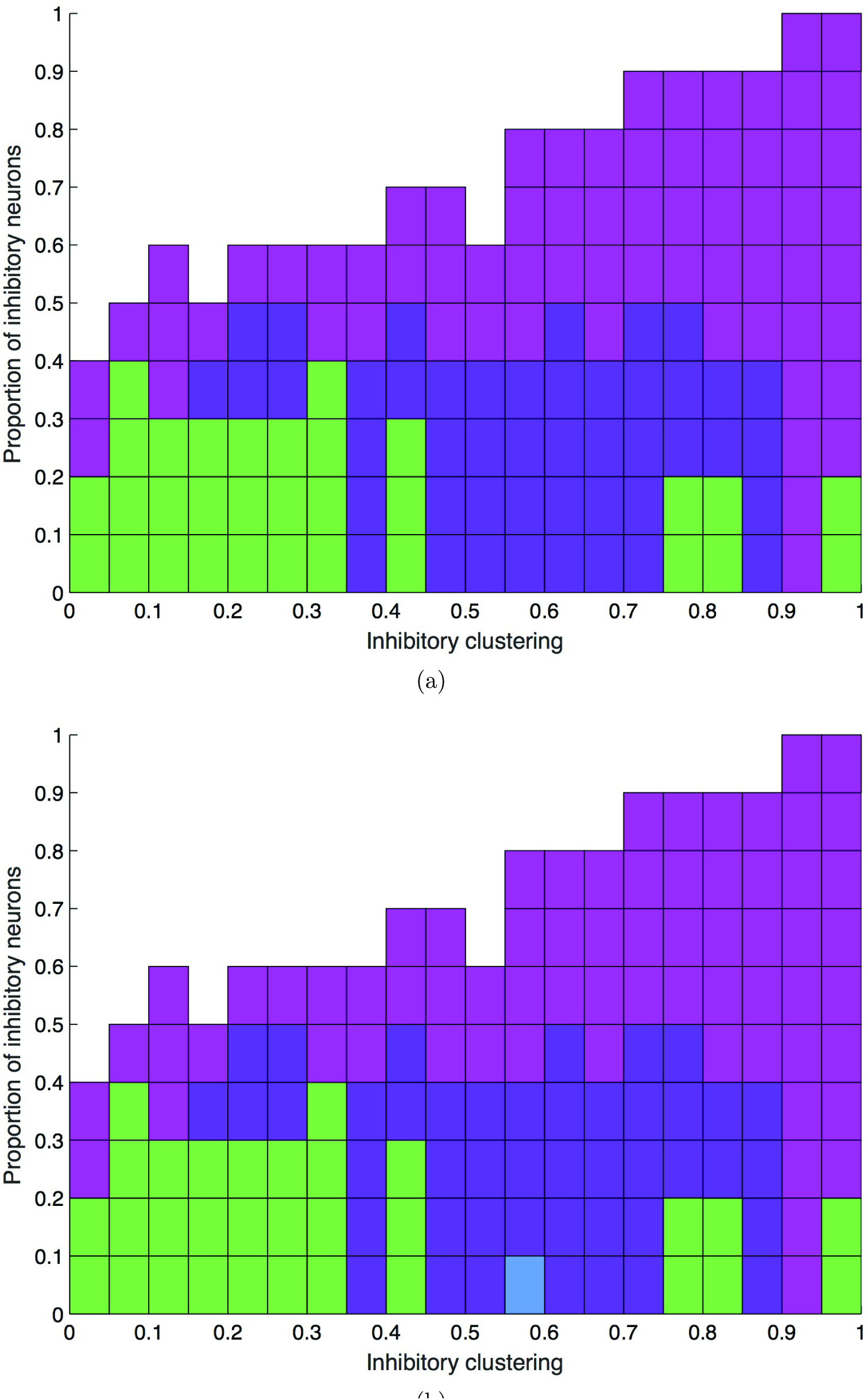
1D system behaviours for growth simulations set-ups with various proportions of inhibitory neurons and levels of inhibitory clustering. Green boxes denote stabilisation of network structure and electrical activity. Purple boxes denote large unstable oscillations in network structure and unstable burst-like oscillations in electrical activity. Blue boxes denote stable oscillatory behaviour of neuritic radii and membrane potentials. Pink boxes denote unbounded growth examples, where the system was unable to reach the desired level of electrical activity for all neurons, with all neuritic fields growing indefinitely. White areas did not have any suitable configurations to run. In panel (a), the networks were initially disconnected. In panel (b), the networks started as regular ring networks.

### 3.3 Is there an optimal proportion of inhibition for inducing oscillatory behaviour in networks containing inhibitory clusters?

When high levels of inhibitory clustering are considered and networks are made up of repeated tile arrangements, thus creating network structures with one or more inhibitory clusters, a proportion of inhibitory neurons in the range [20%, 50%] induced oscillatory behaviour in both network structure and electrical activity (Fig. 5). Within this range, some configurations experienced stable oscillations of network structure and electrical activity with small amplitudes, whilst the majority of other simulations experienced sustained unstable burst-like oscillatory behaviour of electrical activity and sustained unstable oscillations of neuritic radii values with larger amplitudes. For proportions of inhibition below 20%, all simulations experienced complete stabilisation of network structure and electrical activity. With a proportion of inhibition above 50%, the systems were unable to achieve the desired level of electrical activity for every neuron (∊ = 0.6). Neurons which are unable to achieve a membrane potential of ∊ experience unbounded growth of their neuritic field, with their membrane potential stabilising at a level below the homeostatic set-point. In such a situation, a stable network end state is unachievable. We see very similar regimes emerge when the starting network is either totally disconnected (Fig. 5a) or sparsely connected (Fig. 5b). In both cases, we observe a transition between stabilisation and oscillatory behaviours as the proportion of inhibition exceeds 20%, and between oscillatory behaviours and unbounded growth as the proportion of inhibitory neurons exceeds that of excitatory neurons.

**Figure 5:**
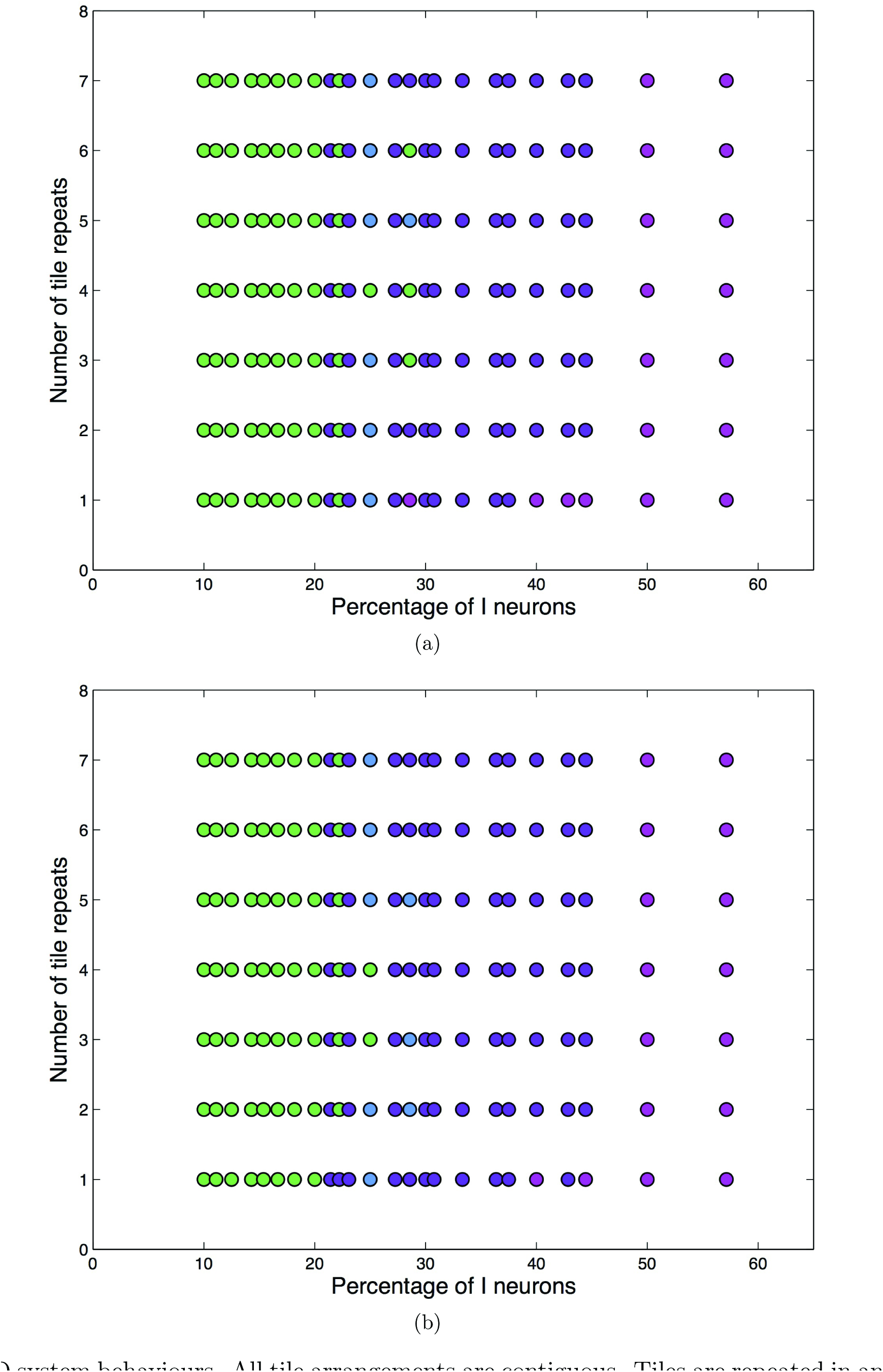
1D system behaviours. All tile arrangements are contiguous. Tiles are repeated in an adjacent manner to maintain a 1D system. Green circles denote networks experiencing complete stabilisation. Blue circles denote periodic (stable) oscillatory behaviour. Purple circles denote unstable oscillatory behaviour. Pink circles denote simulations where all neuritic fields grow unbounded because the desired level of electrical activity cannot be reached. In panel (a), the networks were initially disconnected. In panel (b), the networks started with sparse connectivity, with neuritic radii values randomly generated in the interval (0, 0.6) (identical seed of value 10 for all networks). All other parameters were as listed in Table 1.

### 3.4 Network analysis of global behaviour types

To gain a deeper understanding of how distinct global network behaviours are characterised in terms of network theory measures, we plotted neuritic radii and membrane potential dynamics, alongside corresponding network structure visualisations and degree distributions, for three simulation configurations with distinct global behaviours (Figs. 6-8). Using the subgraph counting algorithm described in Ritchie et al. (2014), we performed motif analysis on adjacency matrices at specific time points from each simulation to determine if global behaviour types are also characterised by specific network motifs (Table 3). Here, we list the behaviour types observed and describe each in terms of structural and electrical dynamics, and we discuss other characteristics of note.

**Figure 6:**
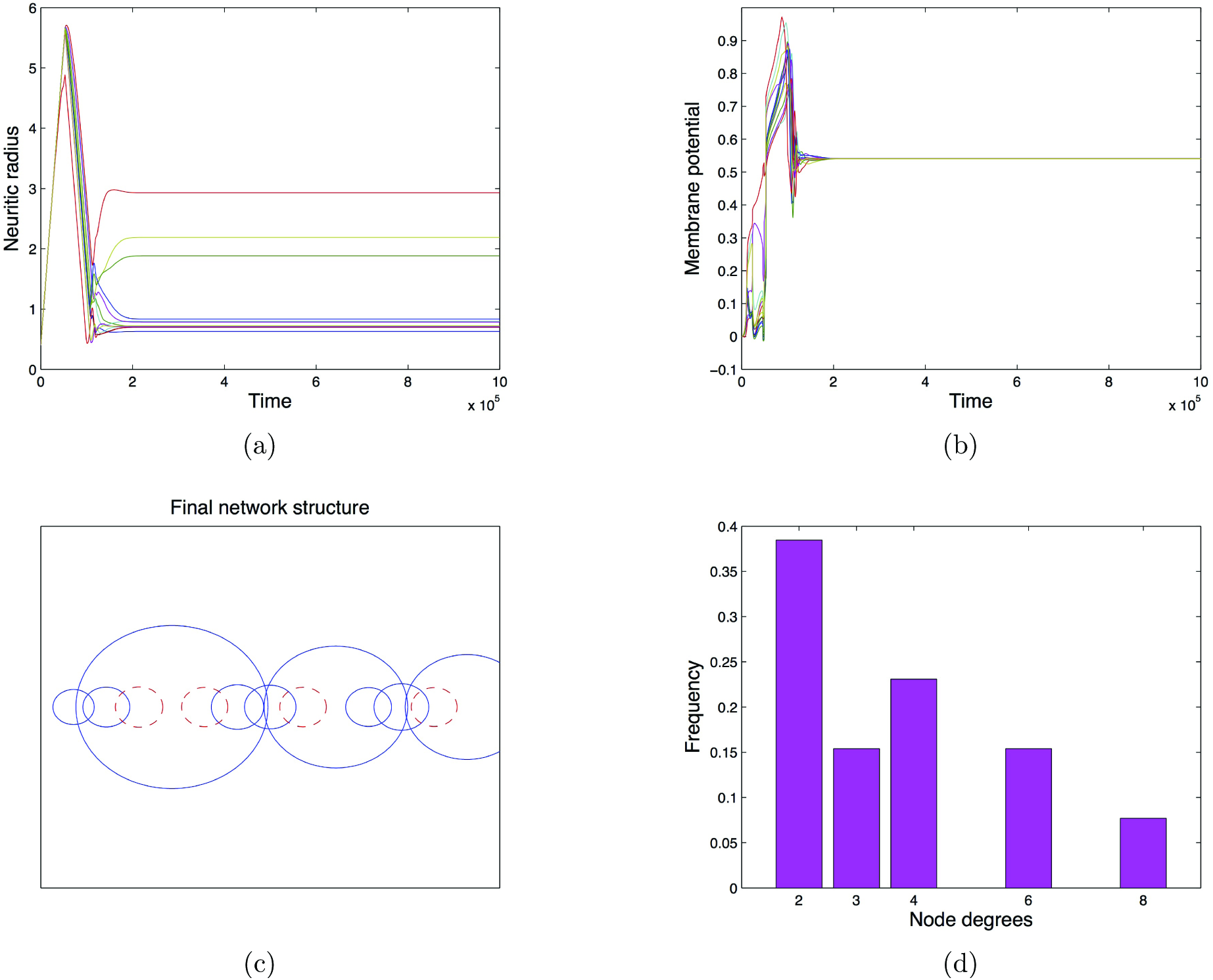
Global system behaviour and analysis of the associated network structure at equilibrium. This is the same simulation as shown in Figs. 1e and 1f, with the spatial arrangement ∘ ∘ • ∘ • ∘ ∘ • ∘ ∘ ∘ • ∘. Panel (a) depicts the dynamics of the neuritic radii, (b) depicts the dynamics of the membrane potential values, (c) is a visualisation of the network’s structure at the end of the simulation, where blue circles denote the field of an excitatory neuron and dashed, red circles denote the field of an inhibitory neuron. Panel (d) shows the degree distribution of all neurons at equilibrium.

**Figure 7:**
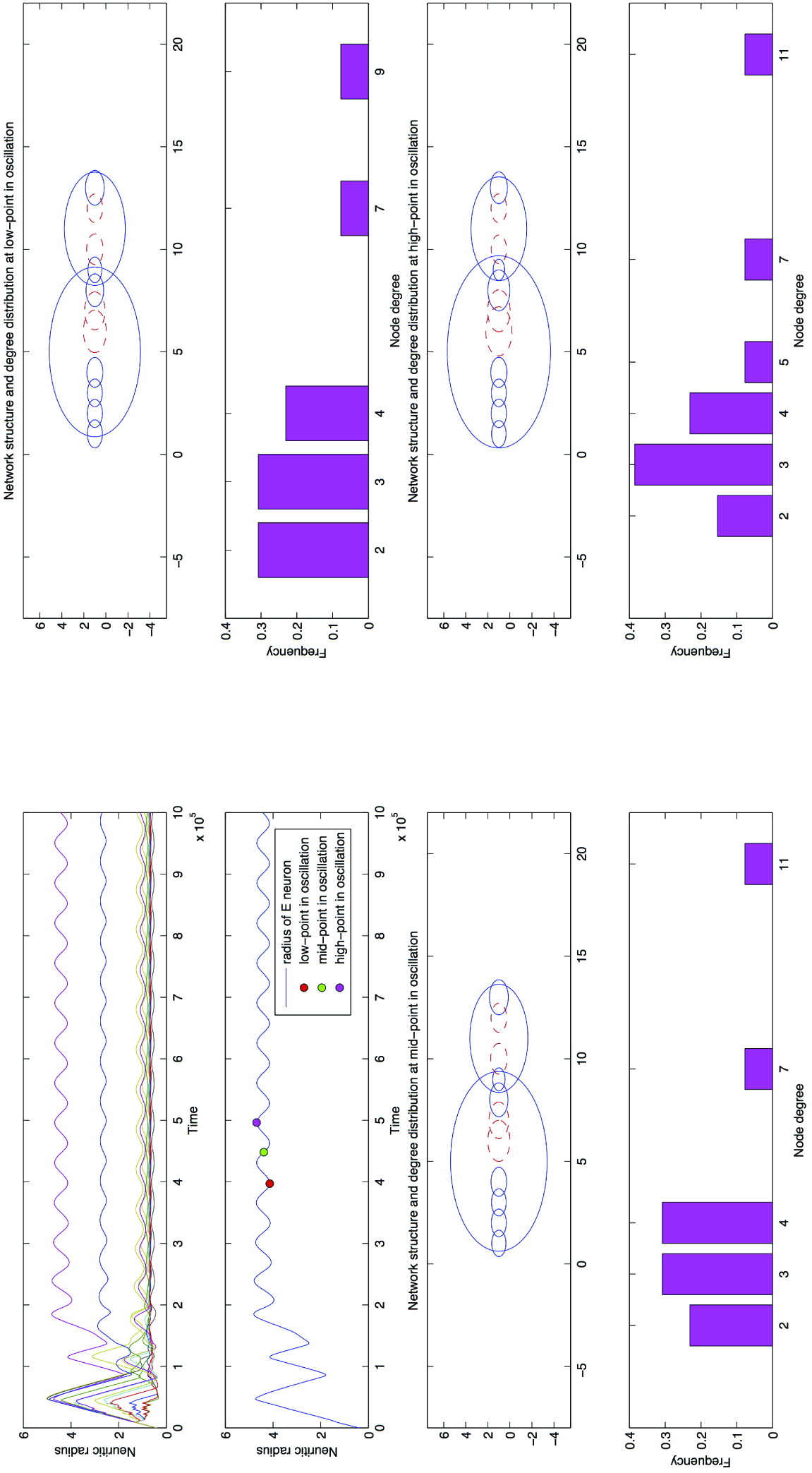
Network structure and degree distribution of the simulation shown in Figs. 1a and 1b, at three time points during the periodic oscillations - see coloured markers in the first two panels, top-left quadrant. The spatial arrangement of neurons is ∘∘∘∘∘••∘∘•∘•∘ with the network initially disconnected. Bottom-left plots show the network structure and associated degree distribution at the low-point of an oscillation, top-right plots show the network structure and associated degree distribution at the mid-point of an oscillation and bottom-right plots show the network structure and associated degree distribution at the high-point of an oscillation. In the network structure plots, solid blue lines denote excitatory neurons and dashed red lines denote inhibitory neurons.

**Figure 8:**
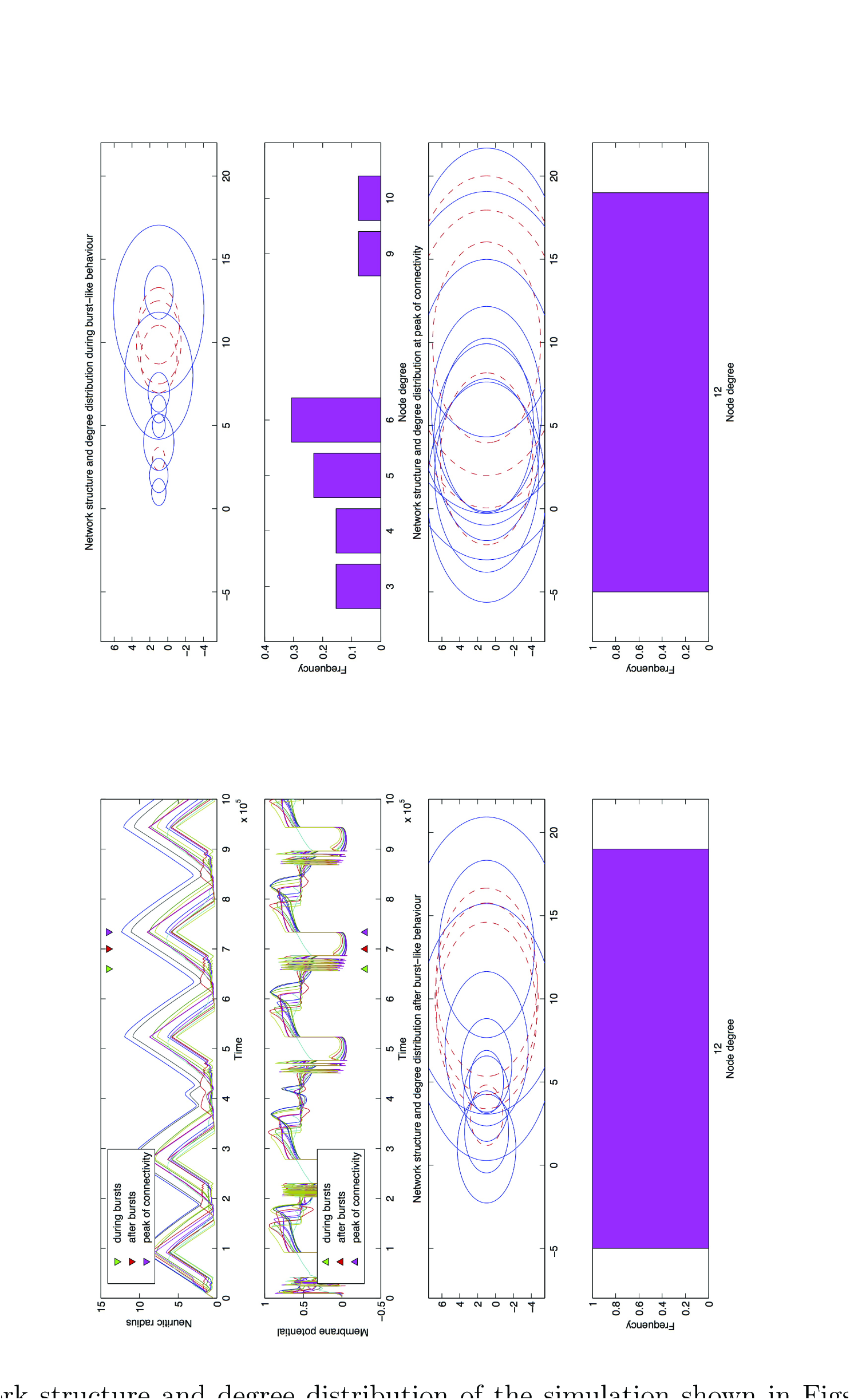
Network structure and degree distribution of the simulation shown in Figs. 1c and 1d, at three time points during the unstable oscillations - see coloured markers in the first two panels, top-left quadrant. The spatial arrangement of neurons is ∘∘∘∘∘••∘∘•∘•∘ with the network initially disconnected. Top-right plots show the network structure and associated degree distribution during the burst-like behaviour. The bottom-left plots depict the network structure and associated degree distribution after the burst-like behaviour and the bottom-right plots show the network structure and associated degree distribution at the peak of connectivity. In the network structure plots, solid blue lines denote excitatory neurons and dashed red lines denote inhibitory neurons.

**Table 3.**
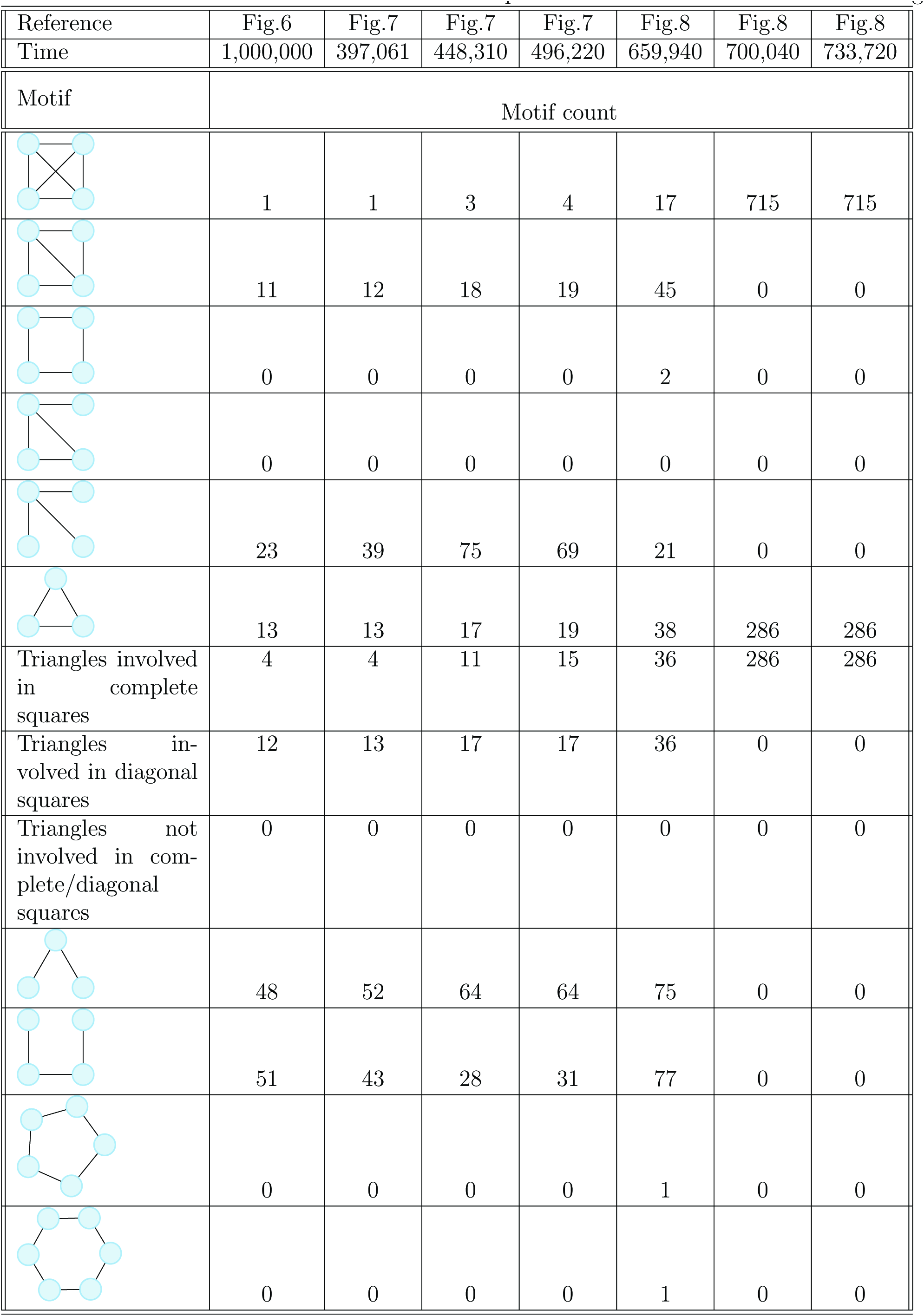
Network motif counts at various points of the simulations shown in Figures 6-8.

#### 3.4.1 Total stabilisation

Networks which experience total stabilisation of their structure and electrical activity (Fig. 6) undergo a period of rapid overshoot in the number of connections, until a critical level of network connectivity is reached (Fig. 6a). At this stage, a rapid pruning of connections occurs, with all neuritic radii shrinking at a similar rate. Simultaneously, we observe an overshoot followed by a steep reduction in the membrane potential for all neurons (Fig. 6b). Following periods of overshoot and pruning, the neuritic radius and membrane potential for all neurons tend quickly towards their equilibrium values. Once all values have stabilised, we look to a visual representation of the final network structure (Fig. 6c) and the corresponding degree distribution (Fig. 6d) to understand the structural elements in more detail. In areas containing inhibitory neurons, neighbouring excitatory neurons have large field sizes, and areas of high inhibitory clustering seem to coincide with even larger excitatory neighbours. The corresponding degree distribution suggests that the introduction of inhibitory neurons has resulted in a long-tailed degree distribution, with the presence of excitatory hub nodes with high degree that are able to contact the majority of the remaining neurons in the system. We note that the inhibitory clustering measure for the network structure (∘∘•∘∘•∘∘∘•∘), which experienced complete stabilisation of both network structure and electrical activity, is 0.04707 (4 s.f). Looking at the resulting network in terms of network motifs present within the structure (Table 3), we observe that the majority of motifs occurring contain relatively few nodes (neurons), e.g. triangular and square motifs.

#### 3.4.2 Stable oscillations

Networks developing stable oscillatory behaviour appear to experience relatively minor changes in their structure over time (Fig. 7). The associated membrane potential dynamics for this simulation (Fig. 1b) depict very similar behaviour to that of the neuritic radii values. After the initial period of overshoot and pruning of network connections, the stable oscillatory regime is reached, and both network structure and electrical activity experience stable oscillations of similar, small amplitudes. Once the system has reached the oscillatory regime, the network structure has a long-tailed degree distribution, indicating the existence of hub nodes. Looking at the dynamics of the two largest neuritic radii (both excitatory neurons neighbouring one or more inhibitory neurons), we observe that their oscillations are almost out of phase with one another; it appears that it could be the interaction between these two large neuritic fields which holds the system in a stable oscillatory regime. The inhibitory clustering measure for the network structure (∘∘∘∘∘••∘∘•∘•∘) is 0.2269 (4 s.f). We note that the network structure during the low-point in the oscillation of one neuritic radius (Fig. 7) contains very similar motifs to the final network structure of the simulation which experienced complete stabilisation (Fig. 6), but by the mid and high-points in the oscillations, the majority of motifs present have increased in number (Table 3).

#### 3.4.3 Unstable oscillations

Networks experiencing sustained unstable oscillatory behaviour undergo far more extreme changes in their structure, level of connectivity and electrical activity over time (Fig. 8). From the start of the simulation, all neuritic radii endure unstable oscillations with large amplitudes, mostly in synchronisation with one another (i.e. all neuritic fields in the system are expanding or retracting simultaneously). During each increase in field sizes, we see bursting behaviour of the membrane potential values, followed immediately afterwards by a return to the resting membrane potential (equal to zero in the ES model). Coinciding with the peak of each oscillation in field size, we see a rapid increase in electrical activity for the majority of neurons. Following this, we observe unstable movements of the membrane potential values and the neuritic fields reduce in size, before the bursting behaviour commences once again. In terms of network attributes during unstable burst-like oscillations, it is clear that the network structure alternates between full and partial connectivity at various points during the oscillations. During the bursting behaviour of the electrical activity, the network structure is not fully connected, with a long-tailed degree distribution that implies the existence of excitatory hub nodes. Some time after the bursting behaviour, the network structure becomes fully connected once again and the cycle of behaviour continues. The inhibitory clustering measure for the network structure (∘∘•∘∘∘∘∘•••∘∘) is 0.2914 (4 s.f). Looking at motif analysis of the network structure at various time points (Table 3), it is immediately obvious that the changes in number and distribution of network motifs during the time-course of this simulation are much more extreme, compared with other behaviour types we analysed in this section. At time 659940, when the membrane potential values are experiencing burst-like behaviour (Fig. 8), the network contains a wide array of motifs, from triangles and squares to pentagons and even hexagons. However, once the network becomes fully connected (after the bursting behaviour and at the peak of network connectivity), we see that the network is entirely made up of fully connected squares.

#### 3.4.4 Unbounded growth

In some network configurations, it is possible for a single neuron not to be able to reach the desired level of activity ϵ. In such case, its neuritic field keeps expanding. However, once the neuritic field overlaps the neuritic fields of all remaining neurons, further increases in field size no longer yield any extra input. Hence, whilst the membrane potential stabilises at a level below the homeostatic set-point, the growth behaviour continues. As mentioned previously, this behaviour is an idiosyncrasy of the model; it is not biologically relevant and therefore does not warrant further analysis.

## 4 Discussion

Previous studies of the ES model have postulated that outgrowth and interactions between excitation and inhibition can lead to complicated patterns of development in individual cells (Van Ooyen et al., 1995), since excitation increases activity but inhibits outgrowth, whereas inhibition does the opposite. When dealing with a purely excitatory model, with individual neurons assigned various outgrowth rates (the parameter *ρ* in the model), complex periodic behaviour of connectivity and electrical activity was observed, with the authors concluding that the precise behaviour depends on the spatial distribution of the cells and the distribution of the outgrowth properties over the cells (Van Ooyen and Van Pelt, 1996). A later study with a simple two-neuron E-I model (van Oss and Van Ooyen, 1997) showed that the introduction of inhibition could account for bistability. It concluded that purely excitatory networks end up in the same attractor regardless of initial conditions, whereas mixed (excitatory and inhibitory) networks do not necessarily do so. Further it shed some light on the excitation inhibition balance by showing that networks developing under conditions of relatively high inhibition end up in a ‘pathological’ state with oscillatory activity whereas a low level of inhibition during the initial stage of development enable the system to move to the ‘normal’ state (van Oss and Van Ooyen, 1997).

Our work reinforces the idea that networks with higher levels of inhibition are more likely to experience oscillatory dynamics of and on the network. However, through revealing distinct global behaviours for different spatial arrangements with a fixed number of excitatory and inhibitory neurons, we have established that the global behavioural dynamics and structural characteristics of the network rely not only on the balance between excitation and inhibition, but also on the spatial arrangement of neurons within the network (and, to a limited extent, the initial conditions). In particular, we have offered that inter-inhibitory connectivity, which we have characterised through a measure of inhibitory clustering, is an important determinant, with networks with higher levels of inhibitory clustering more likely to undergo oscillatory dynamics in electrical activity and network structure than networks with low levels of inhibitory clustering.

There is little published literature available on the effects of increased inhibitory clustering on neuronal network structure and dynamics. In studies of the ES model, inhibitory neurons are found to: impose a structure on neighbouring excitatory neurons, help connect to different parts of a structure by inducing outgrowth, and increase the overall degree of connectivity in a network (Van Ooyen et al., 1995). It was proposed that the number and distribution of inhibitory neurons is important: when inhibitory neurons are able to contact one another (i.e. our clustering measure (8) becomes larger), they are electrically inhibited (self-inhibition), but their outgrowth will become stimulated. This causes the ultimate level of inhibition in the system to become higher than without self-inhibition. It was also noted that long-range inhibition is obtained when inhibitory cells occur in a clustered fashion and are able to stimulate one another’s outgrowth (Van Ooyen et al., 1995). More recently, Litwin-Kumar and Doiron (2012) investigated the implications of *excitatory* clustering on cortical activity, finding that even modest clustering substantially changed the behaviour of the network activity, introducing slow dynamics during which clusters of neurons experience temporary changes in their firing rate. In a network structure consisting of solely excitatory and inhibitory neurons, excitatory and inhibitory clustering will be somewhat congruous to one another, and, in this respect, our results support and complement those of Litwin-Kumar and Doiron (2012).

To test our hypothesis that network structures with high levels of inhibitory clustering were more likely to undergo oscillatory behaviour, we proposed a measure of inhibitory clustering within tile networks, where neurons are assigned to lattice positions. This measure captures the extent of inhibitory clustering within a tile network, and provides ease of comparison across tile network structures of different sizes; there is scope for improvement, however. The measure, see Eq. (8), takes into account the closeness of *M* −1 inhibitory pairs with *M* inhibitory neurons, but does not measure the order of these pairs. Using the current measure, it is therefore possible to find distinct spatial arrangements which produce an identical clustering measure but experience different global behaviours. For example, rows 21 and 22 in Table 2 have the same clustering measure, but one network experiences complete stabilisation whilst the other undergoes stable oscillations. We also note that the inhibitory clustering measure is only valid for ‘one-dimensional’ tile structures; a similar measure for two-and three-dimensional lattice structures is required in order to compare clustering levels in networks of higher dimension.

A simulation of a network tile with a clustering measure of exactly 1 (i.e., containing a contiguous arrangement of neurons) revealed a particular dynamic behaviour, whereby the system appears to transition between regular, small-amplitude stable oscillations of membrane potential and neuritic radii, and less regular burst-like oscillations of electrical activity and network structure with a much larger amplitude (Figs. 3a, 3b). It is unclear what mechanism caused this transition, but Haider et al. (2006) suggested that the dynamic balance between excitation and inhibition may allow for rapid transitions between relatively stable network states.

The inhibitory clustering measure allowed us to map out the relationship between inhibitory clustering and proportion of inhibition in a system, and the associated dynamics of and on the circular tile networks. Irrespective of whether the networks were initially connected or disconnected, we observed a clear structure (see Figs. 4a and 4b respectively). However, it is important to note that the different regimes are extremely course-grained; for each segment plotted, we ran one individual simulation, where the proportion of neurons was contained within an interval of width 10% and the clustering measure was contained within an interval of length 0.05. In order to obtain sufficient coverage of all segments, we considered various network sizes, a range of values for *M* and*N*, and various spatial arrangements. We therefore expect there to be increased variability on the boundaries of the regime areas, and we treat the regimes produced with caution, focussing more on the qualitative results than on specific behaviours.

A similar structure is observed again when plotting the proportion of inhibitory neurons in a tile versus the number of adjacent tile repeats combined to form the entire network (Fig. 5). The idea of having adjacent tile repeats is a similar concept to that explored by Malagarriga et al. (2015) in which mixed cortical macrocolumns operating in a partially synchronised, irregular regime are seen to affect the dynamics of the network itself. The authors find that in this type of mesoscopic network, excitation and inhibition spontaneously segregate, with some columns acting mainly in an excitatory manner while some others have predominantly an inhibitory effect on their neighbours. They also report that hub nodes (i.e. nodes with high degree) are preferentially inhibitory, contrasting with our finding that increased inhibitory clustering induces excitatory neurons of high degree. Whilst we do not suggest that excitatory hubs are physiologically likely, it is worth mentioning that in our networks, hubs are an emerging property whereas in the above study hubs are the result of imposing a scale-free network structure on the model. Further, in agreement with Malagarriga et al. (2015), when we analysed the characteristics of neuronal networks experiencing distinct global behaviours, we observed some emerging segregation of excitation and inhibition, suggesting that clustered inhibitory neurons induce a secondary network ‘layer’ of high-degree excitatory neurons which could be responsible for the transport of information across the whole network. We acknowledge that the networks studied here are biologically unrealistic due to their size and therefore there are limited conclusions to be drawn regarding the presence of network motifs and degree distributions. Network analysis on simulations with larger networks consisting of two and three-dimensional lattices with periodic boundary conditions would provide more insight into the emergence of specific network motifs and their function.

The ES model uses circular neuritic fields to model the area of influence of a single neuron. One consequence of this is that all neurons connect to their direct neighbours before connecting to neurons located further away topologically. As a result, the networks cannot develop the small-world properties observed in neuronal networks. Future work could investigate the effects of inhibitory clustering using an activity-dependent growth model where neurons have a variable and discrete number of synaptic elements, and various synapse formation rules can be implemented (e.g. random, distance-dependent or otherwise), in order to create specific network attributes. Such a modelling approach was shown to conserve small-world network properties in a study investigating cortical reorganisation after focal retinal lesions (Butz and van Ooyen, 2013).

Simulations involving the ES model suggest that the complex dynamical behaviour neuronal networks experience is influenced by a combination of factors, including the balance between excitation and inhibition, the spatial topology of those excitatory and inhibitory neurons, and the connections between them. There are too many intrinsic limitations to the model to be allowed to make strong statements about cortical development. Nevertheless, this work suggests that modelling studies should be careful to attribute particular dynamic properties to the E/I ratio as such properties may exist in well-mixed populations but not in the kind of structured networks found in physiological systems. This certainly highlights the need for further work in developing mathematical models on structured networks.

## Acknowledgements

Rosanna Barnard gratefully acknowledges EPSRC (Engineering and Physical Sciences Research Council) for funding her PhD studentship (Doctoral Training Grants EP/L505109/1 and EP/M506667/1), and the Leverhulme Trade Charities Trust for the awarding of a postgraduate bursary to support her research. We would also like to thank Simon F. Farmer for his helpful comments on an early draft.

